# The differential spatiotemporal expression pattern of shelterin genes throughout lifespan

**DOI:** 10.1101/104299

**Authors:** Kay-Dietrich Wagner, Yilin Ying, Waiian Leong, Jie Jiang, Xuefei Hu, Yi Chen, Jean-François Michiels, Yiming Lu, Eric Gilson, Nicole Wagner, Jing Ye

## Abstract

Shelterin forms the core complex of telomere proteins and plays critical roles in protecting telomeres against unwanted activation of the DNA damage response and in maintaining telomere length homeostasis. Although shelterin expression is believed to be ubiquitous for stabilization of chromosomal ends, some evidence suggests that some shelterin subunits have tissue-specific functions. However, very little is known regarding how shelterin subunit gene expression is regulated during development and aging. Using two different animal models, the mouse and zebrafish, we reveal herein that shelterin subunits exhibit distinct spatial and temporal expression patterns that do not correlate with the proliferative status of the organ systems examined. Together, this work shows that the shelterin subunits exhibit distinct spatiotemporal expression patterns, suggesting important tissue-specific functions during development throughout the lifespan.

## Introduction

Telomeres are specialized chromatin structures, which cap chromosome ends and provide chromosome stability. The maintenance of telomeres requires accurate protections against DNA damage response (DDR) that would otherwise permanently stop cell division by checkpoint activation [ataxia telangiectasia mutated (ATM), and ATM- and Rad3-related (ATR) signaling] and lead to end-to-end chromosomal fusions by non-homologous end joining (NHEJ). Another threat to genome integrity stems from the inability of the conventional replication machinery to fully replicate the extremities of parental DNA, erosion compensated for by telomerase or recombination mechanisms (1,2).

To achieve chromosome end protection, telomeres are composed of repetitive DNA sequences that can fold into a terminal loop (t-loop), nucleosomes, the non-coding telomeric repeat-containing RNA (TERRA), the protein complex shelterin, and an ill-defined network of nuclear factors (3). Shelterin is essential for telomere protection and is composed of six subunits: three bind specifically to telomeric DNA (TRF1, TRF2, and POT1) and three establish protein–protein contacts: RAP1 with TRF2, TIN2 with TRF1 and TRF2, and TPP1 with TIN2 and POT1. Each shelterin subunit appears to have a specific role in telomere protection, i.e., TRF2 blocks ATM signaling and NHEJ, while POT1 blocks ATR signaling (4).

Importantly, telomeres are dynamic structures during development, cancer and aging (5-7). Indeed, the expression of telomerase is repressed in somatic tissues, leading to a progressive and cumulative telomere shortening with cell division ultimately leading to critically short telomeres triggering DDR and cellular senescence (8). Therefore, telomeres have emerged as a key driver of aging.

In addition to their role in chromosome end protection, shelterin subunits are able to localize outside telomeric regions, where they can regulate the transcription of genes involved in metabolism, immunity and neurogenesis (9). This delineates a signaling pathway by which telomeric changes (i.e. telomere shortening) control the ability of their associated factors to regulate transcription throughout the nucleus. This coupling between telomere protection, and tissue-specific transcriptional control might reflect the necessity of tissue homeostasis to rely on ‘fine-tuned’ coordination between telomeric dynamic (reflecting replicative history and the cumulative effects of various types of stress affecting telomere structure), cellular senescence and differentiation (9).

However, although many molecular and animal studies have manipulated the expression levels of shelterin subunits, only a few have examined the organ specificity of shelterin protein expression. We measured here shelterin gene expression levels in various tissues during development and early adulthood of mice and throughout the lifespan of zebrafish. This revealed distinct spatiotemporal regulation patterns of shelterin subunits during development and aging.

## Materials and Methods

All animals were used in accordance with the guidelines of the French Coordination Committee on Cancer Research and local Home Office regulations.

More specific details on the experimental procedures used for immunohistology and qRTPCRs are given in the Supplemental Experimental Procedures. Data are expressed as means ± SEMs. ANOVA together with the Bonferroni post-hoc or Mann-Whitney test was performed as indicated. A p value < 0.05 was considered to reflect statistical significance.

## Results

### Shelterin genes are differentially expressed in mouse tissues

We measured the mRNA levels of mouse shelterin genes (*TERF1, TERF2, RAP1, TPP1, TINF2, POT1a,* and *POT1b*) in mouse brains, hearts, livers, and kidneys, commencing on embryonic day 10 (E10), at 2-day intervals up to postnatal day 1 (P1), and then on P7, P21, and P100 (Figure 1). The proliferating cell nuclear antigen (*PCNA*) gene served as a marker of proliferation in each organ system. In young mice, the relative expression levels of shelterin genes differed among the four tissues evaluated (Figure 1 and Table I). *TPP1* was most prominently expressed in all tissues; whereas *TERF2* and *RAP1* were more highly expressed than was *TINF2* and *POT1a/b* in the brain but not the heart, liver, or kidney. This differential expression pattern appeared to be established during development. Indeed, we observed downregulation of *POT1a*/*b*, unchanged expression of *TINF2* and *TERF1*, and a significant increase in *TERF2, RAP1,* and *TPP1* expression through to adulthood (Figure 1, Table II). All shelterin components except *POT1a* were expressed at low levels in the heart, and the *POT1a* level decreased significantly during adulthood. The levels of *TPP1, RAP1*, and *TERF1* were higher in the liver than in the heart, and *POT1a/b* expression levels decreased during development. Kidney *TPP1* and *RAP1* levels were upregulated during adulthood; *POT1a/b* expression decreased, but no significant changes in the levels of *TERF1*, *TERF2*, or *TINF2* were observed (Figure 1). As expected, *PCNA* gene expression decreased significantly through to adulthood in all organs evaluated, indicating that the expression levels of the various shelterin genes were not associated with the proliferative status of the various tissues.

**Figure 1.**
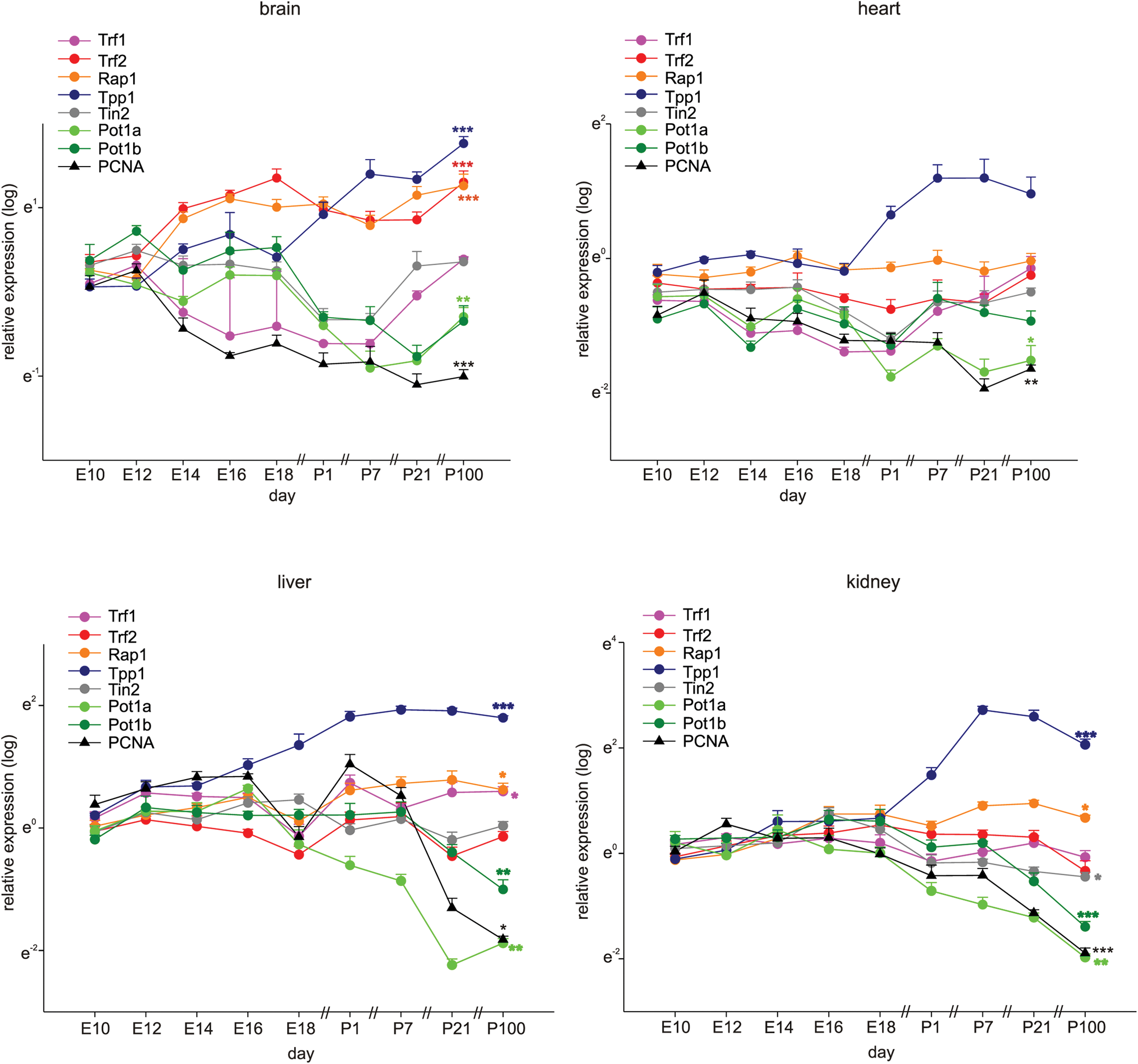
Shelterin components are differentially expressed during development and adulthood. Quantitative RT-PCRs for Shelterin components Trf1, Trf2, Rap1, Tpp1, Tin2, Pot1a, and Pot1b and PCNA as a marker for proliferation in mouse brains, hearts, livers, and kidneys at different time-points of development and in adulthood (*n*=4 each, the four samples for E10 were each pooled from 7 organs, at E12 and 14 the four samples were pooled from four organs each). Significance was tested between E10 and P100 (adult). Data are mean ± SEM. *p<0.05, **p<0.01, ***p<0.001.

**Table I.**
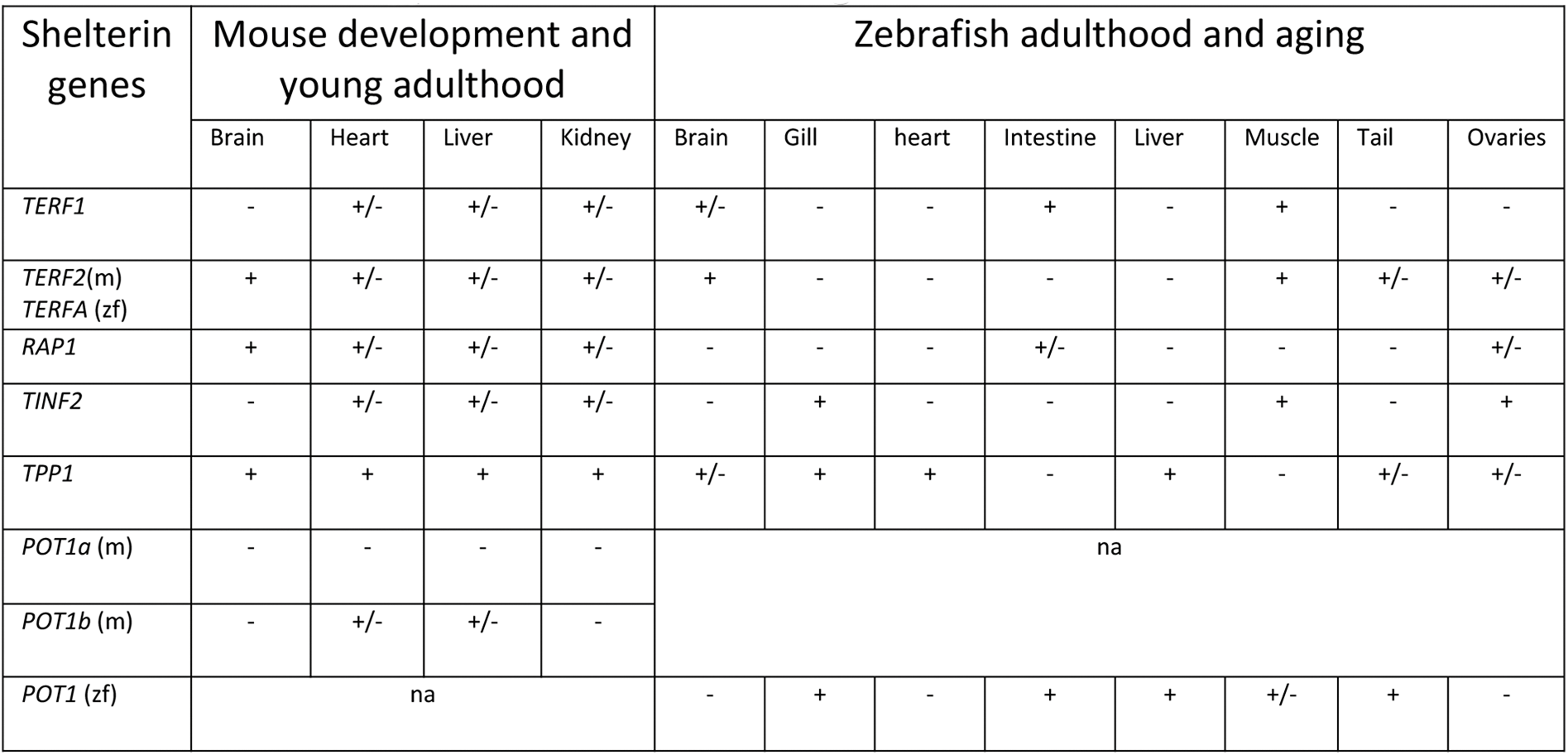
Relative expression of shelterin genes

**Table II.** Trend in shelterin gene expression during development, adulthood and aging

| Shelterin genes | Mouse development and young adulthood |  |  |  | Zebrafish adulthood and aging |  |  |  |  |  |  |  |
| --- | --- | --- | --- | --- | --- | --- | --- | --- | --- | --- | --- | --- |
|  | Brain | Heart | Liver | Kidney | Brain | Gill | heart | Intestine | Liver | Muscle | Tail | Ovaries |
| *TERF1* | s | s | UP | s | DOWN | DOWN | s | DOWN | s | s | DOWN | DOWN |
| *TERF2*(m)<br>*TERFA* (zf) | UP | s | s | s | DOWN | DOWN | s | DOWN | s | DOWN | DOWN | DOWN |
| *RAP1* | UP | s | UP | UP | DOWN | DOWN | s | DOWN | DOWN | DOWN | DOWN | DOWN |
| *TINF2* | s | s | s | s | DOWN | DOWN | DOWN | s | UP | DOWN | DOWN | DOWN |
| *TPP1* | UP | s | UP | UP | DOWN | DOWN | DOWN | s | DOWN | DOWN | DOWN | DOWN |
| *POT1a* (m) | DOWN | DOWN | DOWN | DOWN | na |  |  |  |  |  |  |  |
| *POT1b* (m) | DOWN | s | DOWN | DOWN |  |  |  |  |  |  |  |  |
| *POT1* (zf) | na |  |  |  | DOWN | s | DOWN | DOWN | DOWN | s | DOWN | DOWN |

### TRF2 is highly expressed in the mouse neuronal system during both development and adulthood

To evaluate in detail differential TRF2 protein expression during development, we stained mouse tissues for TRF2 during development until young adulthood (i.e., from E10 to P100). TRF2 was highly expressed in all tissues until E16, at which time expression began to decrease in the heart, liver, and kidney but remained high in the brain (Figure 2). This finding is in contrast to what was reported by Cheng et al. (10), who found that TRF2 was not detected in the brain to E18. This may be attributable to technical problems, as the authors used a mouse-derived antibody to examine mouse tissues. In the absence of extensive blocking procedures, this may create false-negative results caused by an enhanced background. During later development and young adulthood, we found that TRF2 expression decreased in the heart, liver, and kidney but remained stable in the brain; the protein was highly expressed in neurons (Figure 3, Supplementary Figure 1). In the heart, TRF2 continued to be expressed in some endothelial cells of the subepicardial vessels, but in the kidney, the expression thereof became restricted to glomerular podocytes and juxtaglomerular cells (Figure 3), in agreement with the recently described angiogenic properties of the protein and its expression regulation by the Wilms’ tumor suppressor WT1 (11).

**Figure 2.**
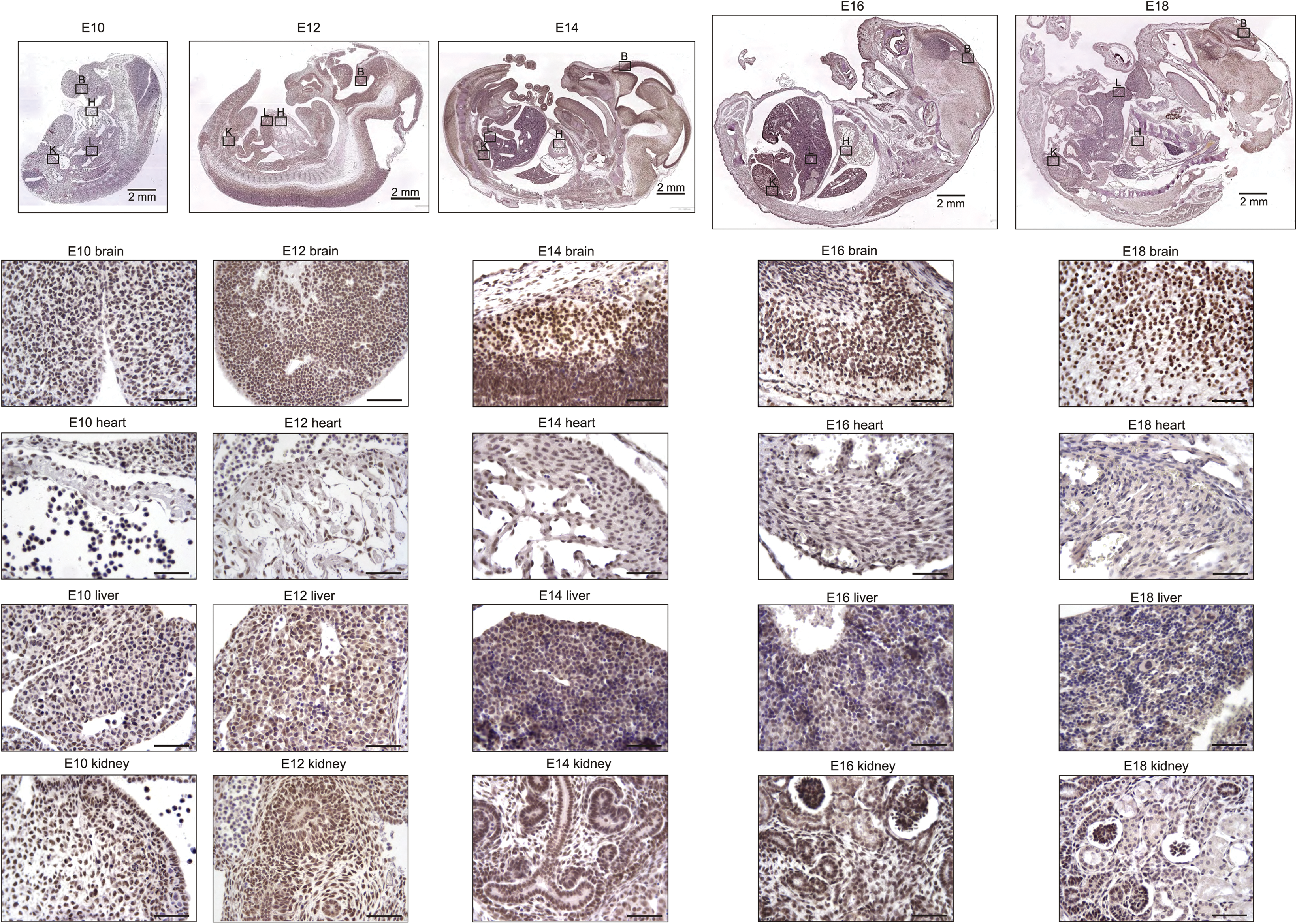
TRF2 is Highly and Ubiquitously Expressed During Embryonic Development Up to E16 and Persists Afterwards Specifically in the Brain. Representative photomicrographs of TRF2 immunostaining on sections of mouse embryos (3,3′ diaminobenzidine (DAB) substrate, brown, hematoxylin counterstaining) at different stages before birth. B: brain, H: heart, L: liver, K: kidney. Unless otherwise indicated, scale bars represent 50µm.

**Figure 3.**
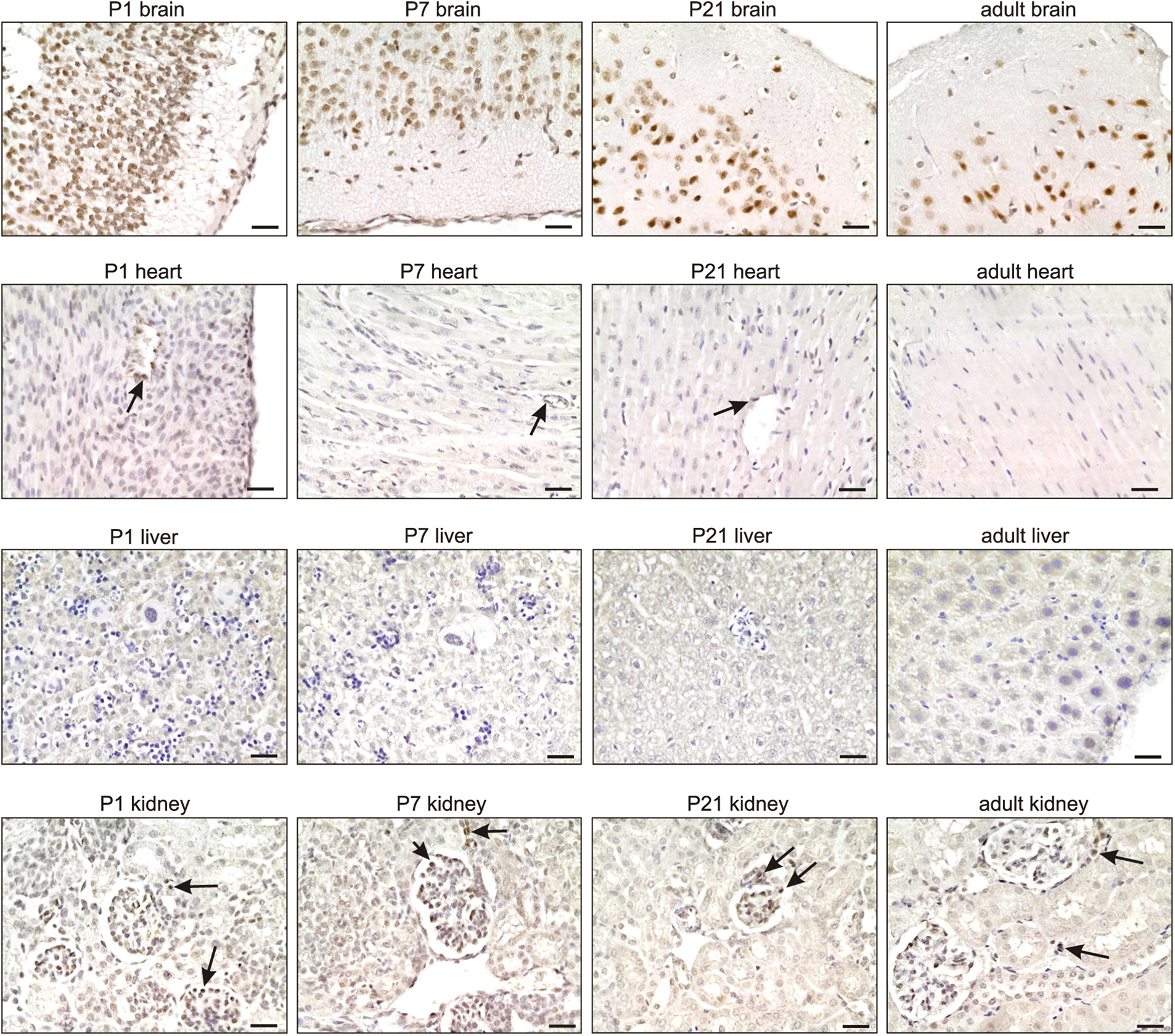
TRF2 Expression Remains High in the Brain During Adulthood. Representative photomicrographs of TRF2 immunostaining for the brain, heart, liver, and kidney (3,3′ diaminobenzidine (DAB) substrate, brown, hematoxylin counterstaining) at different stages after birth. Note the persistent high expression of TRF2 in neurons of the brain (see also Figure S3), the specific expression in subepicardial endothelial cells of the heart, and glomerular podocytes and juxta-glomerular cells of the kidney (arrows). Scale bars indicate 50µm.

The persistent high level of TRF2 expression in the brain during both development and adulthood is in line with previous work showing that TRF2 expression specifically increases upon neural differentiation (12) (10,13). Such expression was accompanied by production of a brain-specific cytoplasmic form of TRF2, termed TRF2-S, which lacks both the DNA-binding domain and the nuclear localization signal (14) (15). However, we failed to detect marked cytoplasmic staining of TRF2 in neurons; the staining was predominantly nuclear (Figures 2-3). Overall, the results suggest that the nuclear form of TRF2 plays a key role in brain development and function.

### Shelterin genes are differentially expressed in zebrafish tissues throughout lifespan

The subunit composition of zebrafish shelterin is similar to that of humans; the complex is composed of the six subunits TRF1, TRF2 (termed TRFA in zebrafish), RAP1, TIN2, TPP1, and POT1. We determined the relevant mRNA levels in various tissues of 6 female fishes, from the young adult stage (3 months) to aged fish (36 months) (Figure 4). We confirmed tissue identities using specific markers (Supplementary Figure 2). As in the mouse (Figure 1), the relative expression levels of shelterin genes varied among tissues. Thus, the relative levels of *TERFA* mRNA were highest in brain and muscle and lowest in liver; *TPP1* mRNA showed highest expression in the heart and lowest expression in the intestine and ovaries (Figure 4, Table I). During aging, we observed a trend toward general downregulation of shelterin gene expression (Figure 4, Table II); this was particularly marked in the brain and ovaries. The relative shelterin gene expression pattern was usually preserved, with the exception of the *RAP1*, which decreased in mRNA expression more rapidly than did the other shelterin genes in the intestine and the gill.

**Figure 4.**
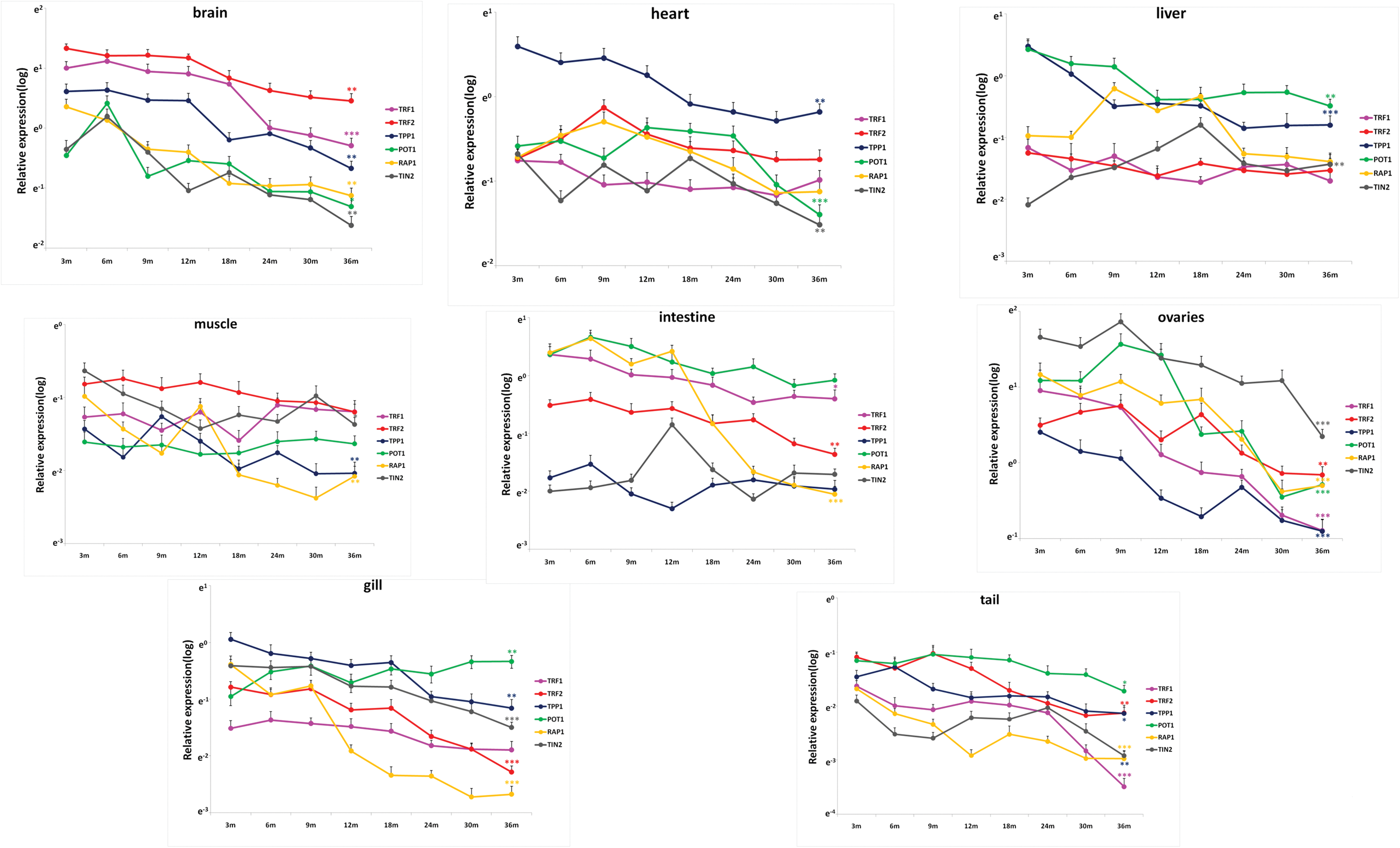
Shelterin Genes are Differentially Expressed during Zebrafish Life Span. Quantitative RT-qPCRs for Shelterin components TRF1, TRF2 (TERFA), RAP1, TPP1, TIN2 and POT1 in zebrafish’s brain, heart, liver, intestine, muscle, gill, tail and ovary at different time-points of life span from 3 month to 36 month (n==6 each). Significance was tested between 3 month and 36 month. Data are mean ± SEM. *p<0.05, **p<0.01, ***p<0.001.

We performed whole-mount *in situ* hybridization of zebrafish embryos for *TERFA* mRNA using a cDNA probe (Figure 5). The signal corresponding to *TERFA* mRNA was present throughout the entire embryo from the blastula (4 hpf) to the gastrula (8 hpf) stage. In contrast, the neuronal marker, Neurog1 (16), was detected only in neuronal tissues and only from 12 hpf to hatching at 72 hpf, whereas the hematopoiesis factor c-MYB was detected only in hematopoietic tissue and then only during the late stages of development (17)(Figure 5). Interestingly, high expression of *TERFA* mRNA in the nervous system was maintained from the beginning of the somite stage to the time of hindbrain formation (20 hpf) and thereafter (Figure 5). As was also true of Neurog1, at 20 hpf, *TERFA* mRNA appeared to be expressed prominently in the dorsal root ganglion and midbrain boundary, in the regions of the neural tube that give rise to the neocortex, midbrain, and hindbrain, in the dorsal and ventral spinal cord, and in regions of the peripheral nervous system. These results are in agreement with the quantitative reverse-transcription PCR (qRT-PCR) data on zebrafish tissues (Figure 4) and the specific neuronal staining of TRF2 during mouse development (Figures 2–3 and supplementary Figure 2). In summary, TRF2 expression appears to be ubiquitous during early development but becomes progressively more restricted to neuronal tissues during later stages of development and into young adulthood.

**Figure 5.**
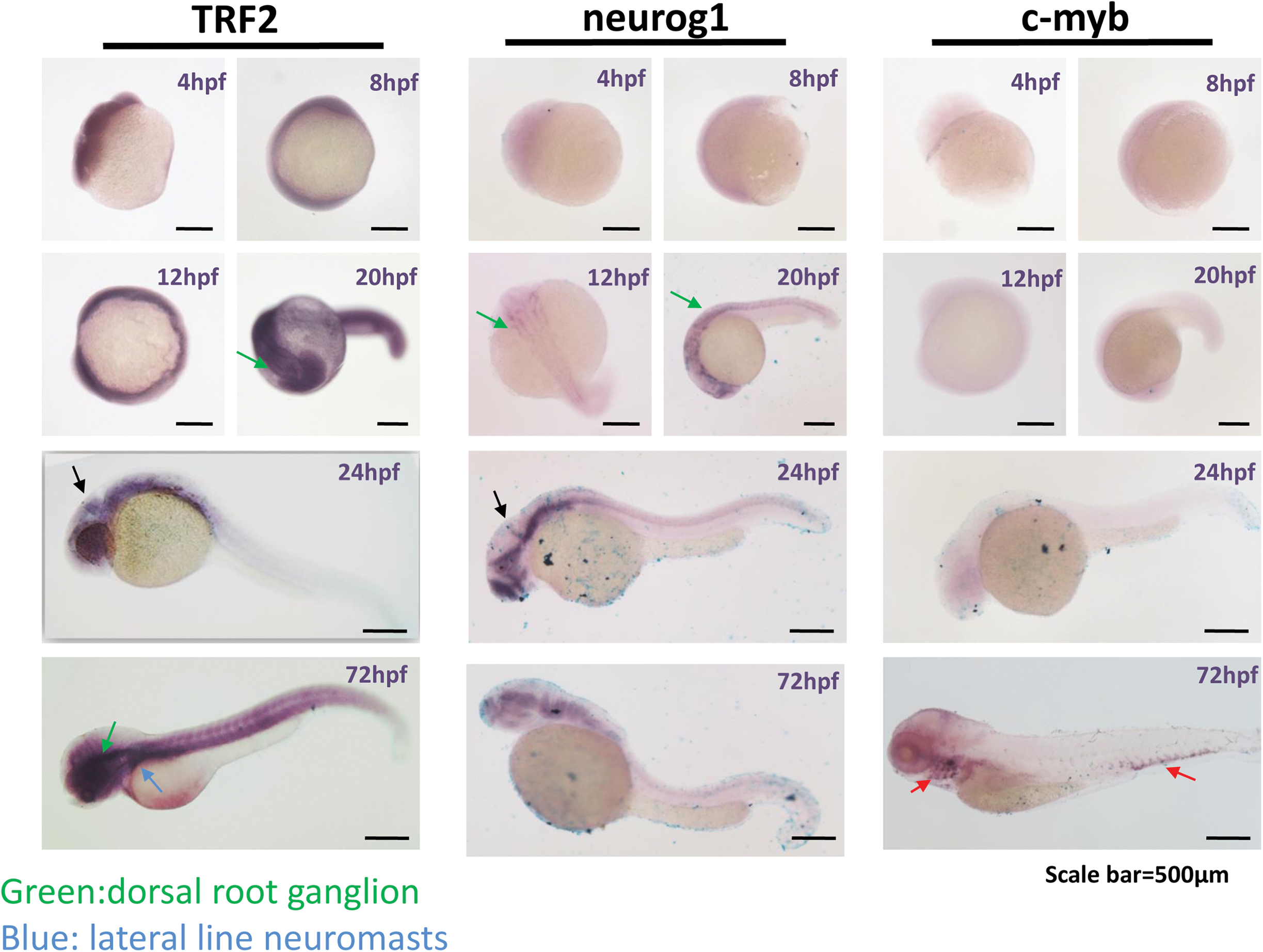
TRF2 (TERFA) Expression Increase since Neural Development at Embryonic Stage and Remains High in the Brain during Larval Stage Development. Representative photomicrographs of whole-mount in situ hybridization of TRF2 (TERFA), Neurog1 and c-MYB. The RNA probe labelled with DIG was stained in dark blue. The green arrow indicates the dorsal root ganglion neuron and the blue arrow indicates the lateral line neuromasts. The black arrow in 24hpf indicates the midbrain boundary. The red arrow indicates the c-MYB signal marked hematopoietic tissue.

## Discussion

We unveil here a tissue-specific shelterin gene expression pattern that is largely conserved between mouse and zebrafish (Table I). In particular, the relative expression level of *TPP1* mRNA was high in most mouse and fish tissues, whereas the *TERF2/TERFA* mRNA levels were specifically elevated in the brain. With the exception of a study on tissue-specific expression of human shelterin genes in response to physiological stress (18), the present study is, to the best of our knowledge, the first to show that shelterin gene expression levels change in a tissue-specific manner during development and aging. We propose that this spatiotemporal expression pattern of shelterin gene expression plays important roles during development, tissue homeostasis, and aging. The fact that TRF2/TRFA is highly expressed in the brain, a tissue of low proliferative activity, indicates that the tissue-specific roles played by TRF2 (and probably other shelterin components) may (at least in part) be independent of the functions in telomere protection.

Our study had certain limitations. First, the tissues evaluated are composed of many different cell types, and thus we cannot clearly conclude whether the changes noted are suggests that the more proliferative cell types do not significantly express shelterin genes. Second, most of our significant findings were detected at the mRNA level. However, we found that the TRF2 spatiotemporal expression pattern was identical, both, on the mRNA and the protein level, during mouse development, suggesting that any effects of posttranscriptional regulation may be limited. Overall, our results suggest that important tissue-specific shelterin subcomplexes exist. This is consistent with previous studies on the tissue-specific roles played by TRF2 (13,14) and RAP1 (19) (20). Consequently, the mechanisms by which shelterin and telomere structures affect cell fate may be more varied than previously thought; specific shelterin subcomplexes may be associated with different cell fates.

The existence of tissue-specific shelterin gene expression brings to light the view that tissue homeostasis relies on ‘fine-tuning’ of the coordination between the telomeric state and cell-type-specific functions (9). These mechanisms may explain the broad contributions made by telomeres and telomerase to normal development and aging, as well as the roles played by dysregulation of telomeres and telomerase in cancer and various other tissue-specific pathologies (6).

Our data form a solid foundation for future studies exploring the physiological role and regulation of the shelterin complex during development and aging. The work raises several questions. How is the spatiotemporal pattern of shelterin gene expression involved in tissue development, renewal, and function? How is shelterin gene expression regulated during development and over the subsequent lifespan? In this context, we have previously shown that the TERF2 gene is a direct target of Wnt/beta-catenin and WT1 in both mouse and human cells (21) (11), suggesting that these signaling pathways play important roles in the spatiotemporal expression of shelterin during development and aging. If shelterin plays a critical role in telomere protection, how do the extratelomeric functions of shelterin subunits contribute to cell-type-specific functions? Interestingly, a global decrease in shelterin gene expression during aging was evident in most of the tissues evaluated. How is this decrease triggered during aging? Does the decrease actually cause aging, and, if so, can we stop aging by restoring normal levels of shelterin subunits? Future experiments addressing these questions will certainly shed new light on the increasingly complex and dynamic interaction between telomeres and lifespan.

## Acknowledgements

### Author Contributions

NW, KDW, EG, and JY designed the experiments. KDW, YLY, NW, WL, JJ, YC, and XFH performed the experiments. YE, EG, KDW, NW, YLY, and JFM analyzed the data. NW, KDW, EG, and YE wrote the paper. EG, NW and JY are co-senior authors because of their specific roles in coordinating the whole study (EG), the experiments with mice (NW) and the experiments with zebrafishes (JY). JY is the lead author.

### Acknowledgements

Work in the JY/YL laboratories was supported by the National Natural Science Foundation of China (grant numbers 81000875, 81171846, 81270433, 81372099, 81471400, and 81522017), the Shanghai Foundation for Basic Research of Science and Technology, China (grant number 13JC1404001), the Foundation for Committee of Science and Technology in Shanghai (grant number 11ZR1422100), and the Shanghai Municipal Education Commission (Oriental Scholars Program). Work in the EG laboratory was supported by the Ligue Nationale Contre le Cancer (Equipe Labélisée). KDW was supported by the Association pour la Recherche sur le Cancer and Fondation de France. The work was also supported by the French Government (National Research Agency, ANR) through the "Investments for the Future" LABEX SIGNALIFE program (reference ANR-11-LABX-0028-01).

Conflict of interest: none.

